# A Framework for SPICE MRSI Reconstruction Posterior Uncertainty Quantification

**DOI:** 10.64898/2026.11.05.764591

**Authors:** Tian Lyu, Saad Jbabdi, Simon M. Finney, William T. Clarke

**Affiliations:** Oxford Centre for Integrative Neuroimaging, FMRIB, Nuffield Department of Clinical Neurosciences, University of Oxford, Oxford, United Kingdom

**Keywords:** magnetic resonance spectroscopic imaging, low-rank reconstruction, uncertainty quantification, Laplace approximation, posterior covariance, SPICE

## Abstract

**Purpose:** Low-rank reconstruction methods such as SPICE substantially improve Magnetic Resonance Spectroscopic Imaging (MRSI) reconstruction quality. However, SPICE does not provide an estimate of reconstruction uncertainty. A framework for posterior uncertainty estimation in SPICE reconstruction is proposed, hereafter referred to as the framework.

**Theory and Methods:** A Laplace approximation is used to estimate the covariance of the reconstructed spatial coefficients. This covariance is propagated through the spectral basis and spectral fitting to obtain metabolite concentration uncertainty. Matrix-free Hessian–vector products and voxel-wise conjugate-gradient estimation are used to avoid explicit construction or inversion of the full Hessian, making the method tractable.

**Results:** Validation using repeated in vivo 7 T MRSI acquisitions showed that the estimated posterior uncertainty exhibited spatial and spectral patterns similar to those of empirical repeated-scan uncertainty. Experiments further demonstrated how spatial regularization and k-space undersampling affect the estimated uncertainty.

**Conclusion:** The framework provides a practical framework for uncertainty quantification when using SPICE

## Introduction

Magnetic Resonance Spectroscopic Imaging (MRSI) enables spatially resolved measurement of neu-rochemical and metabolic features, providing information about brain chemistry and cellular processes [1]. However, reliable high-resolution MRSI is challenging because of its intrinsically low sensitivity and contamination by nuisance signals from water and lipids that are approximately three orders of magnitude stronger [2].

SPICE (Spectroscopic Imaging by exploiting Spatiospectral CorrElation) is a low-rank MRSI reconstruction framework [3–6] that exploits partial separability [7, 8] and anatomy-informed spatial regularization [9] to enable high-resolution and accelerated reconstruction from undersampled acquisitions. However, SPICE-reconstructed spectra are generated within a predefined spectral sub-space and are therefore noise-free, making the uncertainty of the reconstructed spectra and fitted metabolite concentrations difficult to assess.

Quantifying uncertainty is important because similar estimates may have different levels of reliability, and uncertainty can be incorporated into subsequent statistical analysis [10, 11]. Repeated measurements and bootstrap methods characterise variability across datasets [12], whereas Bayesian posterior uncertainty describes parameter uncertainty conditioned on the acquired data and an assumed model [13], allowing uncertainty to be estimated from a single acquisition. Posterior uncertainty has been applied across quantitative imaging, including quantitative MRI [14, 15], diffusion MRI [16], dynamic contrast-enhanced MRI [17], and PET [18].

However, uncertainty quantification has not been established for the MRSI reconstruction process. We therefore aim to develop a framework for quantifying uncertainty in regularized low-rank MRSI reconstruction, using SPICE [5] as a representative example in this work. In SPICE, uncertainty depends on the measurement noise, Fourier encoding, low-rank spectral basis, and spatial regularization. Direct sampling of the high-dimensional reconstruction posterior is computationally expensive, while explicit evaluation of the covariance is impractical because it involves the inversion of large matrices. An efficient approximation is therefore needed to estimate uncertainty while retaining the structure of the SPICE reconstruction problem.

In this work, we propose a framework for estimating posterior uncertainty in SPICE-reconstructed MRSI data which can be generalised to other regularized low-rank MRSI reconstruction methods. Throughout this work, we use uncertainty to refer to the standard deviation of the posterior distribution. Our framework interprets the SPICE solution as a maximum-a-posteriori (MAP) estimate and uses a Laplace approximation to estimate the covariance of the reconstructed spatial coefficients from the inverse Hessian of the SPICE objective. The covariance of the spatial coefficients is propagated using the spectral basis to obtain “pre-fitting” uncertainty of the image-space MRSI data, and then through spectral fitting to estimate metabolite concentration uncertainty. Matrix-free Hessian–vector products and voxel-wise conjugate-gradient solvers enable the required covariance to be estimated without explicitly constructing or inverting the full Hessian.

We show that the posterior uncertainty obtained through our framework demonstrates strong spatial and spectral agreement with repeated-scan uncertainty. We further investigate how the SPICE spatial regularization parameters and k-space undersampling affect the reconstruction and estimated uncertainty. At present, the implementation is limited to 2D single-slice MRSI, with extension to fully three-dimensional reconstruction remaining as future work.

## Theory

### SPICE Reconstruction Model

Under the low-rank SPICE representation, the MRSI data are modelled as **UV**^*H*^, where **U** ∈ ℂ^*p*×*r*^ contains the spatially varying coefficients and **V** ∈ ℂ^*f* ×*r*^ contains the pre-trained spectral basis functions. Here, *p* is the number of spatial voxels, *r* is the SPICE rank, and *f* is the number of spectral samples. The superscript *T* denotes the ordinary transpose and superscript, *H* denotes the conjugate transpose.

SPICE reconstruction [5, 6, 19, 20] estimates the spatial coefficient matrix **Û** from sampled MRSI *k*–*t* data by jointly enforcing consistency with the acquired data (first term) and an anatomyinformed spatial constraint (second term):

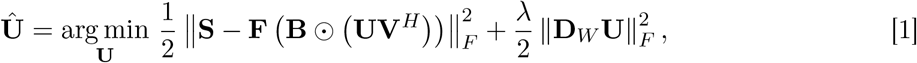

where **S** denotes the sampled *k*–*t* data, **F** is the Fourier sampling operator, **B** represents the spatially dependent *B*_0_ modulation, ⊙ is the Hadamard product, and ∥·∥_*F*_ denotes the Frobenius norm. Since Eq. 1 is quadratic in **U**, the corresponding linear system was solved iteratively using the conjugate-gradient method.

The second term in Eq. 1 imposes an edge-preserving spatial constraint derived from anatomical prior information [4, 5, 9]. Here, **D**_*W*_ is a weighted spatial difference operator that couples neigh-bouring voxels according to their anatomical similarity, while *λ* controls the overall strength of the spatial regularization. The spatial weight is defined as

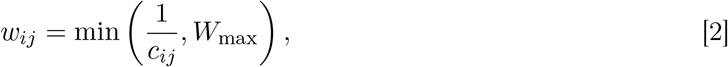

where *c*_*ij*_ is the difference in anatomical-image signal between neighbouring voxels *i* and *j* (i.e., *c*_*ij*_ = |*a*_*i*_ − *a*_*j*_|, where *a*_*i*_ and *a*_*j*_ denote the corresponding anatomical-image intensities) [9]. Voxels with similar anatomical signal therefore have smaller *c*_*ij*_, larger *w*_*ij*_, and stronger spatial smoothing, whereas the weight is reduced across anatomical boundaries. The parameter *W*_max_ caps the weight to prevent 1*/c*_*ij*_ from becoming excessively large when neighbouring anatomical voxels have very similar signal intensities.

In 2D slice implementations, **D**_*W*_ ∈ R^*q*×*p*^ is constructed with one row for each neighbouring voxel pair (*i, j*), where *q* is the total number of such pairs. The corresponding row contains 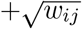 at voxel *i*, 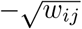 at voxel *j*, and zeros elsewhere, such that the corresponding row of **D**_*W*_ **U** is 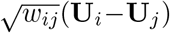. Rows were constructed for vertical, horizontal, diagonal, and anti-diagonal neighbouring pairs [9].

Since the spectral basis represents a low-rank spectral subspace rather than individually identifiable metabolite components, spectral fitting is subsequently performed using FSL-MRS [21] to estimate metabolite concentrations [4, 5, 9].

### Overview of the Proposed Uncertainty Quantification Framework

As illustrated in Fig. 1, our framework extends the SPICE reconstruction pipeline by estimating the posterior covariance of the reconstructed spatial coefficients and propagating it to the reconstructed spectra and fitted metabolite concentrations. In this study, pre-fitting uncertainty is represented by the posterior uncertainty at each voxel and spectral location, while metabolite concentration uncertainty is represented by the posterior standard deviation.

**Figure 1.**
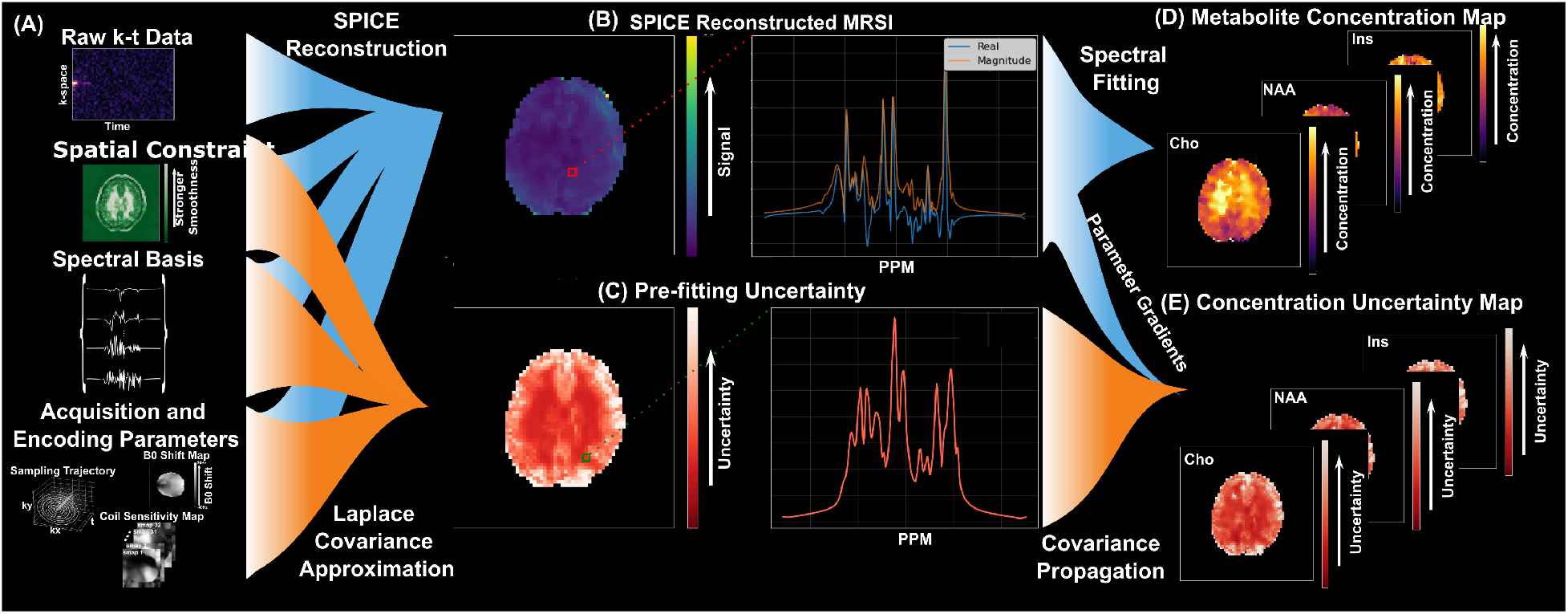
Overall framework for posterior uncertainty estimation in SPICE reconstruction. Blue flow: SPICE reconstruction and spectral fitting pipeline; orange flow: uncertainty estimation before and after spectral fitting. (a) The inputs include the sampled *k*–*t* data, the spatial constraint map derived from the anatomical prior image, the pre-trained spectral basis, and the acquisition and encoding parameters, including the *B*_0_ field map, coil sensitivity maps, and sampling trajectory. (b) SPICE reconstructs the MRSI data by estimating the spatial coefficients from these inputs. (c) Our framework estimates the posterior covariance of the reconstructed spatial coefficients using a Laplace approximation and propagates this covariance through the spectral basis to obtain voxel- and frequency-resolved pre-fitting uncertainty. (d) Spectral fitting is performed on the reconstructed MRSI data to obtain metabolite concentration maps. (e) The pre-fitting covariance is further propagated through the spectral fitting model to obtain metabolite concentration uncertainty.

### Pre-fitting Uncertainty via Laplace Approximation

The SPICE objective function in Eq. 1, which we denote *L*(**U**), contains two terms. Under a Bayesian interpretation, these can be interpreted as a negative log-likelihood term and a prior term, respectively. Minimising *L*(**U**) yields a maximum-a-posteriori (MAP) point estimate. We approximate the posterior distribution *p*(**U** | **S**) with a multivariate Gaussian centred around Û using the Laplace method.

The Laplace approximation describes the local posterior uncertainty using the curvature of the SPICE objective around the MAP estimate. We assume that the noise follows a zero-mean complex Gaussian distribution, *CN* (0, *σ*^2^). The noise variance σ^2^ was estimated from the first 10 noise-only transients of the water-reference acquisition. The complex noise variance was calculated separately for each receiver coil and averaged across the 32 coils to obtain a global noise variance. The posterior covariance is approximated as

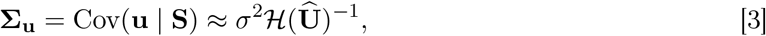

where H(**Û**) denotes the Hessian of the SPICE objective function evaluated at the MAP estimate, and **u** = vec(**U**) is a vectorised version of **U**.

The Hessian matrix can be derived analytically by differentiating the gradient in Eq. 4 with respect to **U**.

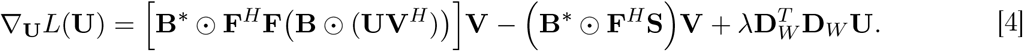

Let **D**_*B*_ = diag(vec(**B**)) and **Φ** = **F**^*H*^**F**. The matrices **I**_*p*_ and **I**_*r*_ denote identity matrices of sizes *p* and *r*, respectively, ^∗^ denotes complex conjugation, and ⊗ denotes the Kronecker product. The overall Hessian evaluated at the MAP estimate is given by:

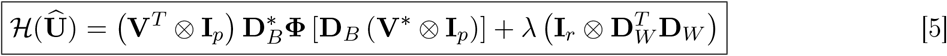

The first term represents the contribution from the data consistency term, and the second term represents the contribution from spatial regularization.

The uncertainty of the reconstructed spectrum can be estimated on a voxel-wise basis. Let **M** ∈ ℝ^*r*×*pr*^ denote a voxel-selection matrix. The multiplication **Mu** extracts the *r* spatial coefficients associated with that voxel. Therefore, the covariance of these coefficients can be obtained from the corresponding block of the inverse Hessian,

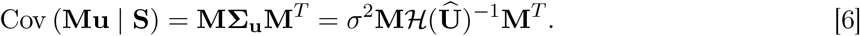

The selected coefficients are then propagated through the spectral basis to obtain the reconstructed spectrum for that voxel,

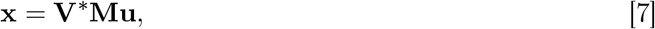

where **x** ∈ ℂ ^*f*^ denotes the reconstructed spectrum of the selected voxel. Since Eq. 7 is a linear transformation of **u**, applying the covariance propagation rule [22] gives

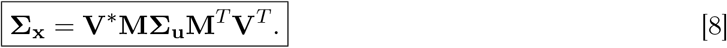

For each voxel, complex spectral samples were drawn from the posterior distribution defined by **Σ**_**x**_ ∈ ℂ *f* ×*f*. The standard deviation across these samples was calculated at each of the *f* spectral points, giving an uncertainty spectrum of length *f* per voxel. The uncertainty spectra from all *p* voxels were then stacked according to their spatial positions to form the final posterior uncertainty map of size *p* × *f*.

### Interpretation of Reconstruction Variance

In the data-consistency term of Eq. 5, (**V**^∗^ ⊗ **I**_*p*_) maps *δ***u** into the spatial–spectral MRSI representation, **D**_*B*_ applies the spatially varying *B*_0_ modulation, and **Φ** = **F**^*H*^**F** propagates it through the sampling pattern, including point-spread-function-induced spatial blurring for non-Cartesian sampling. The adjoint operators then map the response back into the spatial coefficient domain. The regularization term, 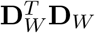, additionally constrains differences between neighbouring coefficients according to anatomical similarity. Stronger overall constraints therefore lead to smaller posterior variance.

### Efficient Voxel-wise Covariance Estimation

According to Eq. 3, direct evaluation of the posterior covariance would require inversion of *H*(**Û**) ∈ ℂ ^*pr*×*pr*^, which is computationally prohibitive for high-resolution MRSI. Instead, we compute only the voxel-specific *r* × *r* block of *H*(**Û**)^−1^ required in Eq. 8.

For a selected voxel, the *r* columns of **M**^*T*^ define the right-hand sides of *r* linear systems with *H*(**Û**). In compact form, *H*(**Û**)**Z** = **M**^*T*^. We solve these systems using conjugate gradient, giving **Z** = *H*(**U**)^−1^**M**^*T*^ without explicitly forming the inverse Hessian.

Each conjugate-gradient iteration requires the action of H(**U**) on a vector. This vector is reshaped as ***η*** ∈ ℂ ^*p*×*r*^, with the same dimensions as **U**, allowing the Hessian action to be evaluated using the original SPICE operators through a matrix-free Hessian–vector product [23]:

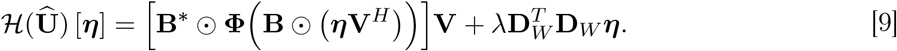

Left multiplication by **M** then extracts the required voxel-specific block, **M***H*(**Û**)^−1^**M**^*T*^ = **MZ**. Although only this *r* × *r* block is retained, it is obtained from linear systems involving the full Hessian and therefore retains the global coupling introduced by Fourier encoding, *B*_0_ correction, the spectral basis, and spatial regularization. The block is scaled by *σ*^2^ according to Eq. 6 and propagated through the spectral basis using Eq. 8.

### Propagation to Metabolite Concentration Uncertainty

The covariance obtained in Eq. 8 characterises the uncertainty of the reconstructed spectrum for each voxel. The metabolite concentrations are subsequently estimated through spectral fitting, which can be regarded as a nonlinear mapping

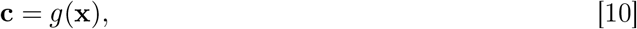

where *g*(·) denotes the spectral fitting model implemented in FSL-MRS [21], and **c** denotes the corresponding metabolite concentration vector. At the reconstructed spectrum **x**, the fitted concentration estimate is 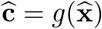.

A first-order Taylor expansion around the reconstructed spectrum 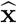 gives 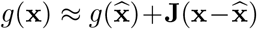 where 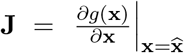 is the Jacobian of the spectral fitting model evaluated at 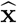. In practice, FSL-MRS provides the analytical Jacobian *∂***x***/∂***c**, for which a small parameter change gives *δ***x** ≈ (*∂***x***/∂***c**)*δ***c**. Recovering the corresponding concentration change from *δ***x** is therefore a local linear least-squares problem, whose solution is *δ***c** ≈ (*∂***x***/∂***c**)^+^*δ***x**. Hence, the Jacobian required for uncertainty propagation is approximated as **J** = *∂***c***/∂***x** ≈ (*∂***x***/∂***c**)^+^, where superscript + denotes the Moore–Penrose pseudoinverse.

According to the first-order uncertainty propagation rule [22],

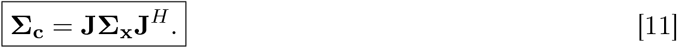

Finally, the posterior standard deviation of each metabolite is obtained from

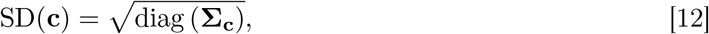

where SD(**c**) contains the posterior uncertainty of the fitted metabolite concentrations.

## Methods

### Data Acquisition

In vivo MRSI data were acquired from two healthy male volunteers using a Siemens MAGNETOM 7T Plus scanner (Siemens Healthineers, Erlangen, Germany), equipped with a Nova Medical Inc. 8Tx/32Rx head coil (Wilmington, MA, USA). One participant underwent five consecutive MRSI acquisitions and was used for the repeated-scan analyses and the representative experiments reported below. The second participant was used to confirm the applicability of the framework on an independent in vivo dataset.

Water suppression and magnetisation preparation used the 2Π-CSAP scheme as described in [24], which employs chemical-shift-selective adiabatic refocusing pulses. Pulse and timing parameters were implemented as described in Supporting Information Table S1 (Non-editing TE=82 ms) of reference [24]. The sequence was implemented in pulseq, using pypulseq [25]. To acquire the water reference, the water suppression module was removed and the frequency-selective band is centred on water’s frequency.

Spatial–spectral encoding was performed using a concentric-ring trajectory comprising 180 rings. The acquisition field of view was 240 × 240 mm^2^ with an in-plane matrix of 64 × 64, corresponding to a nominal in-plane voxel size of 3.75 × 3.75 mm^2^. The slice thickness was 10 mm, resulting in a nominal voxel volume of 3.75 × 3.75 × 10 mm^3^.

The acquisition parameters, identical for the MRSI and water-reference measurements, were TE = 82 ms, TR = 1500 ms, and a flip angle of 15°. The nominal readout duration was 0.4 s, of which approximately the first 0.20 s (198.74 ms) was retained for reconstruction. Each MRSI acquisition lasted 186 s (3 min 06 s) and was repeated five times consecutively, resulting in a total MRSI acquisition time of 930 s (15 min 30 s), excluding the water-reference measurements. The measurement-noise variance *σ*^2^ was estimated from the noise-only transients acquired at the beginning of the water-reference scan.

### Data Reconstruction

The acquired MRSI data were reconstructed using the SPICE framework in Eq. 1, with interchangeable non-Cartesian Fourier encoding backends based on MRI-NUFFT [26] and TorchKbNufft [27]. For TorchKbNufft, repeated evaluation of **Φ** = **F**^*H*^**F** was accelerated using a precomputed Toeplitz kernel [28], applied by FFT-based convolution during reconstruction and covariance estimation.

## Experimental Design

### Comparison of SPICE and Iterative-NUFFT Repeated-Scan Uncertainty

To evaluate the effect of the SPICE reconstruction model on uncertainty, the five repeated acquisitions from the same participant were reconstructed using both SPICE and iterative-NUFFT, and their empirical repeated-scan uncertainties were compared in the spatial and spectral domains.

### Posterior Uncertainty and Repeated-Scan Uncertainty

The framework-quantified posterior uncertainty from a single SPICE reconstruction was compared with the empirical uncertainty across five repeated reconstructions. Although these represent different statistical quantities, their spatial–spectral agreement indicates whether the framework identifies locations that also show greater repeated-scan uncertainty. Agreement was quantified using a single Spearman rank correlation over the pooled voxel–spectral observations within the brain mask.

### Sensitivity to Spatial Regularization Parameters

To investigate the sensitivity of SPICE reconstruction and the uncertainty quantified in the frame-work to spatial regularization, *λ* and *W*_max_ were varied independently, with the other parameter held constant: *W*_max_ = 5 × 10^3^ during the *λ* sweep and *λ* = 1 × 10^−4^ when *W*_max_ was swept. Our method was used to estimate both pre-fitting and metabolite concentration uncertainty for each setting.

For each case, an L-curve was constructed from the data-consistency residual and spatial-regularization norm in Eq. 1 to visualise the trade-off across parameter settings.

### Sensitivity to k-Space Undersampling

Retrospective undersampling was designed following the variable-density strategy commonly used in SPICE, where central k-space is fully sampled and peripheral k-space is more sparsely sampled [5, 29, 30]. Representative SPICE implementations retain approximately the central 30% of the phase-encoding extent and undersample the periphery using undersampling factors of *R* = 2–3. The outermost ring at *k*_max_ was retained in all cases. SPICE reconstruction and uncertainty estimation using the proposed framework were performed for each sampling pattern, and voxel-wise uncertainty ratios relative to the fully sampled reference were summarised across the brain and at a representative voxel.

## Results

### Comparison of Repeated-Scan Uncertainty Between Iterative-NUFFT and SPICE

Fig. 2 compares SPICE and iterative-NUFFT reconstructions and repeated-scan uncertainty. The two methods produced similar overall signal amplitudes (Fig. 2a), although SPICE spectra appeared smoother, with less visible noise, and showed global signal attenuation relative to iterative-NUFFT reconstructions, owing to the applied regularization.

**Figure 2.**
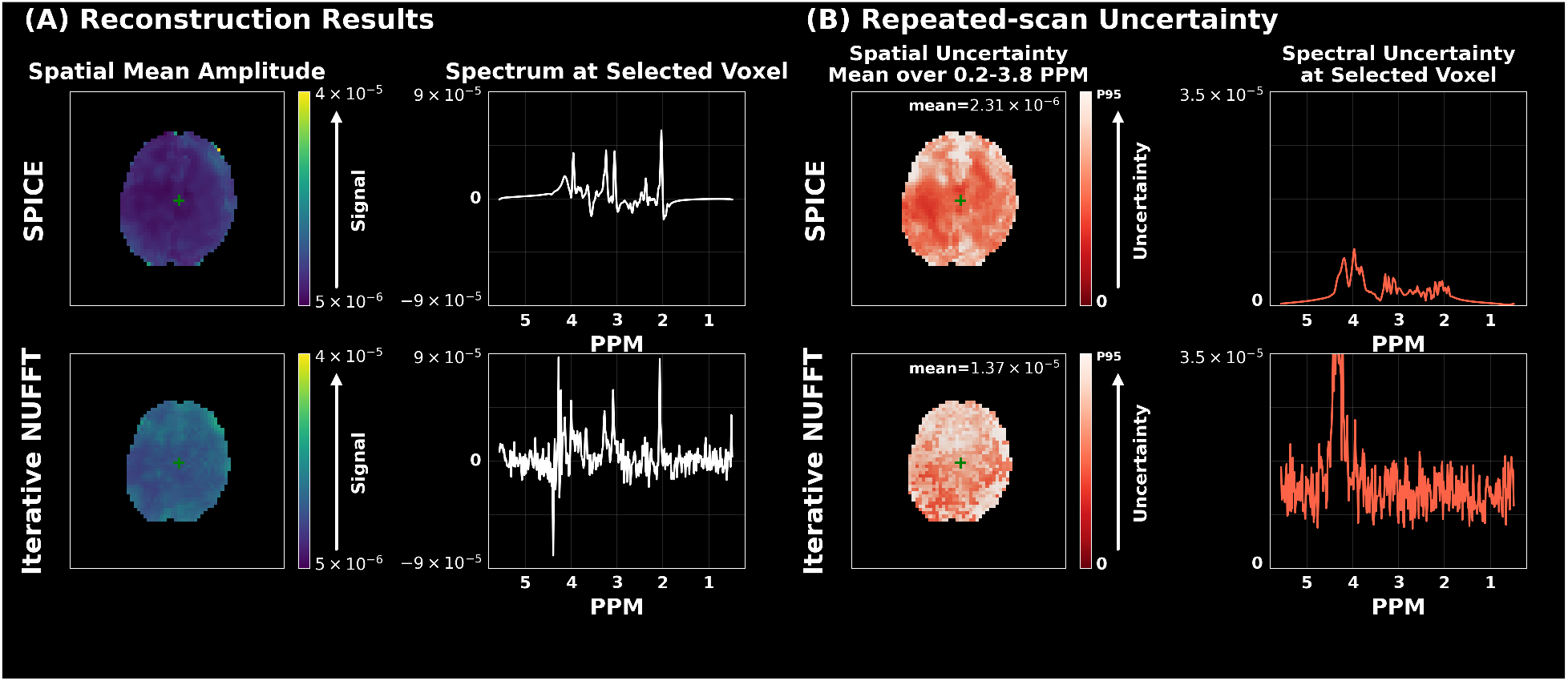
Comparison of reconstruction and repeated-scan uncertainty between SPICE and iterative-NUFFT. (a) Mean FID-amplitude maps and spectra at the indicated voxel reconstructed using SPICE and iterative-NUFFT. Both methods produce similar overall signal amplitudes, while the SPICE spectrum shows substantially lower noise and some signal attenuation due to the low-rank representation and spatial regularization. (b) Repeated-scan uncertainty calculated from five acquisitions for the two reconstruction methods. SPICE shows substantially lower repeated-scan uncertainty with clear spatial and spectral structure, including higher uncertainty near anatomical boundaries and around major metabolite resonances, whereas iterative-NUFFT shows higher and more spatially and spectrally uniform uncertainty outside the region containing residual water.

Repeated-scan uncertainty was markedly lower for SPICE and exhibited clear spatial and spectral structure (Fig. 2b). Higher spectral uncertainty occurred around major metabolite resonances, consistent with variations being constrained to the low-rank spectral subspace. Spatially, uncertainty was higher near anatomical boundaries, where the edge-preserving spatial constraint provides weaker coupling. Additional anterior uncertainty may reflect motion and physiological fluctuations across acquisitions. In contrast, iterative-NUFFT uncertainty was higher and more uniformly distributed because it does not impose the low-rank spectral basis or anatomy-informed spatial constraint. Uncertainty peaks around the water resonance in iterative-NUFFT were associated with variations in residual water signal.

### Posterior Uncertainty and Repeated-Scan Uncertainty

As shown in Fig. 3, the framework-estimated posterior uncertainty from a single SPICE reconstruction and the empirical uncertainty across five repeated acquisitions exhibited similar spatial and spectral patterns. Regions and spectral locations assigned higher uncertainty by the framework showed greater variability across the repeated reconstructions.

**Figure 3.**
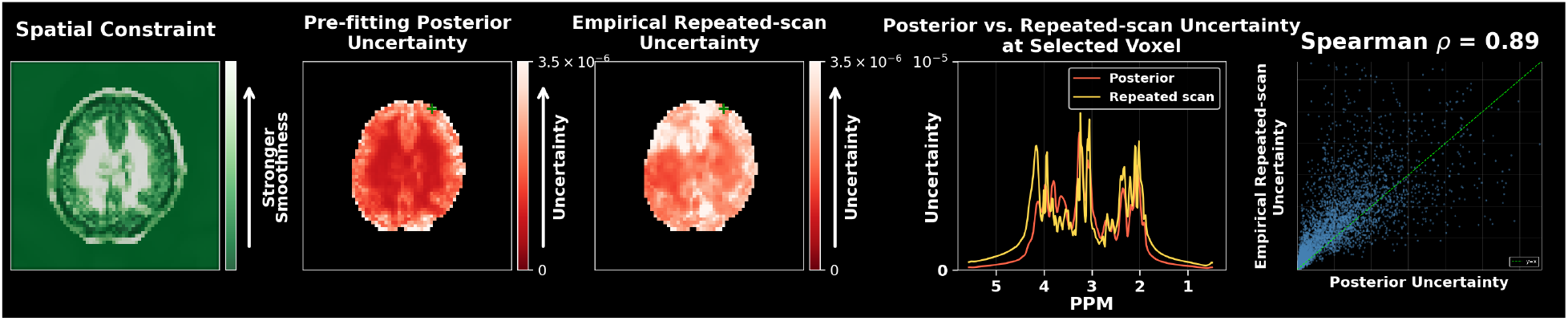
Comparison between framework-estimated pre-fitting posterior uncertainty and empirical repeated-scan uncertainty. Posterior uncertainty estimated using the framework from a single SPICE reconstruction is compared with the empirical uncertainty across five repeated SPICE acquisitions. A representative result obtained using *λ* = 1 × 10^−4^ and *W*_max_ = 5 × 10^3^ is shown. The two measures exhibit similar spatial and spectral patterns, with regions of higher posterior uncertainty generally corresponding to greater repeated-scan uncertainty. Their similar spatial–spectral rank ordering is reflected by a pooled Spearman correlation of *ρ* = 0.89. The two measures are not expected to be numerically identical because repeated-scan uncertainty also includes acquisition-related effects not represented in the current framework.

Posterior uncertainty and empirical repeated-scan uncertainty represent different statistical quantities. Our framework characterises uncertainty conditioned on a single acquisition, the assumed measurement-noise model, and the spatial prior, whereas repeated measurements additionally contain uncertainty from motion, physiological fluctuations, and scanner drift. Similar distinctions between posterior uncertainty and empirical uncertainty have been considered in Bayesian image reconstruction and neuroimaging [18, 31, 32]. Although the posterior and repeated-scan distributions should not be expected to be numerically identical, their spatial and rank-order concordance indicates that uncertainty estimated from an individual acquisition can identify voxels that are more susceptible to variation under repeated measurement.

For each value of *λ*, agreement was quantified using a single Spearman rank correlation over the pooled voxel–spectral observations within the brain mask. The correlation remained high across the nine investigated settings from *λ* = 1 × 10^−6^ to 1 × 10^−2^, with a mean coefficient of *ρ* = 0.9011. For the representative reconstruction at *λ* = 1 × 10^−4^ and *W*_max_ = 5 × 10^3^, the pooled correlation was *ρ* = 0.8850, shown as 0.89 in Fig. 3.

### Effect of the Spatial Regularization Strength λ

Fig. 4 shows the effect of varying the spatial regularization strength *λ*, with *W*_max_ = 5×10^3^ fixed, on estimated concentration and uncertainty maps. Representative results are shown for *λ* = 1 × 10^−6^, 1 × 10^−4^, and 1 × 10^−2^. As *λ* increased, strengthening spatial regularization, the reconstruction appeared progressively smoother (Fig. 4a). However, stronger regularization also increased signal attenuation, as apparent in the spectral domain (Fig. 4b).

**Figure 4.**
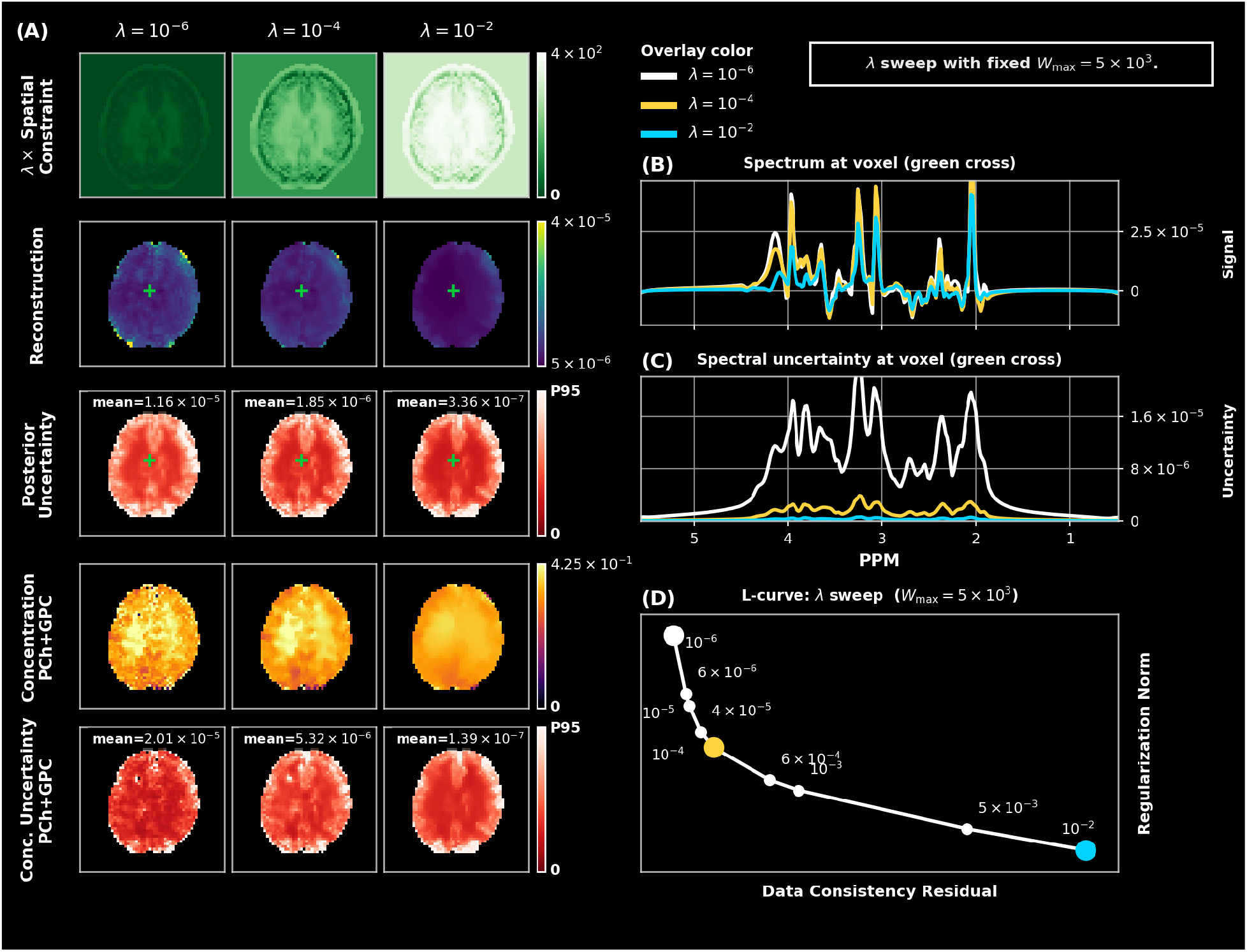
Effect of spatial regularization strength on reconstruction and uncertainty. Results are shown for *λ* = 1 × 10^−6^, 1 × 10^−4^, and 1 × 10^−2^, with *W*_max_ = 5 × 10^3^. The left columns show, from top to bottom, the weighted spatial constraint, reconstructed signal, pre-fitting posterior uncertainty, PCh+GPC concentration maps, and the corresponding concentration uncertainty maps. For visualisation, each pre-fitting uncertainty map and concentration uncertainty map is independently normalised by its within-mask 95th-percentile value (P95), while the annotated mean reports the corresponding unnormalised value. The green crosses indicate the selected voxel used for the overlaid spectra and spectral uncertainty shown on the right. Increasing *λ* produces smoother reconstructed and metabolite maps, reduces spectral fluctuations and signal amplitude, and decreases both pre-fitting posterior uncertainty and metabolite concentration uncertainty. The L-curve summarises the trade-off between the data-consistency residual and regularisation norm over the full *λ* sweep.

Quantified pre-fitting posterior uncertainty decreased with increasing *λ*, became more spatially structured, and had clearer tissue-level contrast (Figs. 4a and 4c). The fitted PCh+GPC concentration maps also became progressively smoother, with substantial loss of local spatial variation at the highest *λ*. In contrast, the corresponding concentration-uncertainty maps showed lower overall variance and a more structured spatial distribution (Fig. 4b). The L-curve summarises the trade-off between data consistency and spatial regularization (Fig. 4d).

### Effect of the Edge-Preserving Constraint Parameter W_max_

Fig. 5 shows the effect of varying *W*_max_ with *λ* fixed. Representative results are shown for *W*_max_ = 1 × 10^2^, 5 × 10^3^, and 1 × 10^5^. Increasing *W*_max_ enforces stronger similarity between anatomically similar neighbouring voxels while retaining a weaker smoothness penalty across anatomical boundaries. Consequently, the reconstructed signal and metabolite concentration maps have sharper tissue boundaries (Fig. 5a). Unlike the *λ* sweep, the selected-voxel spectra showed comparatively little signal attenuation (Fig. 5b).

**Figure 5.**
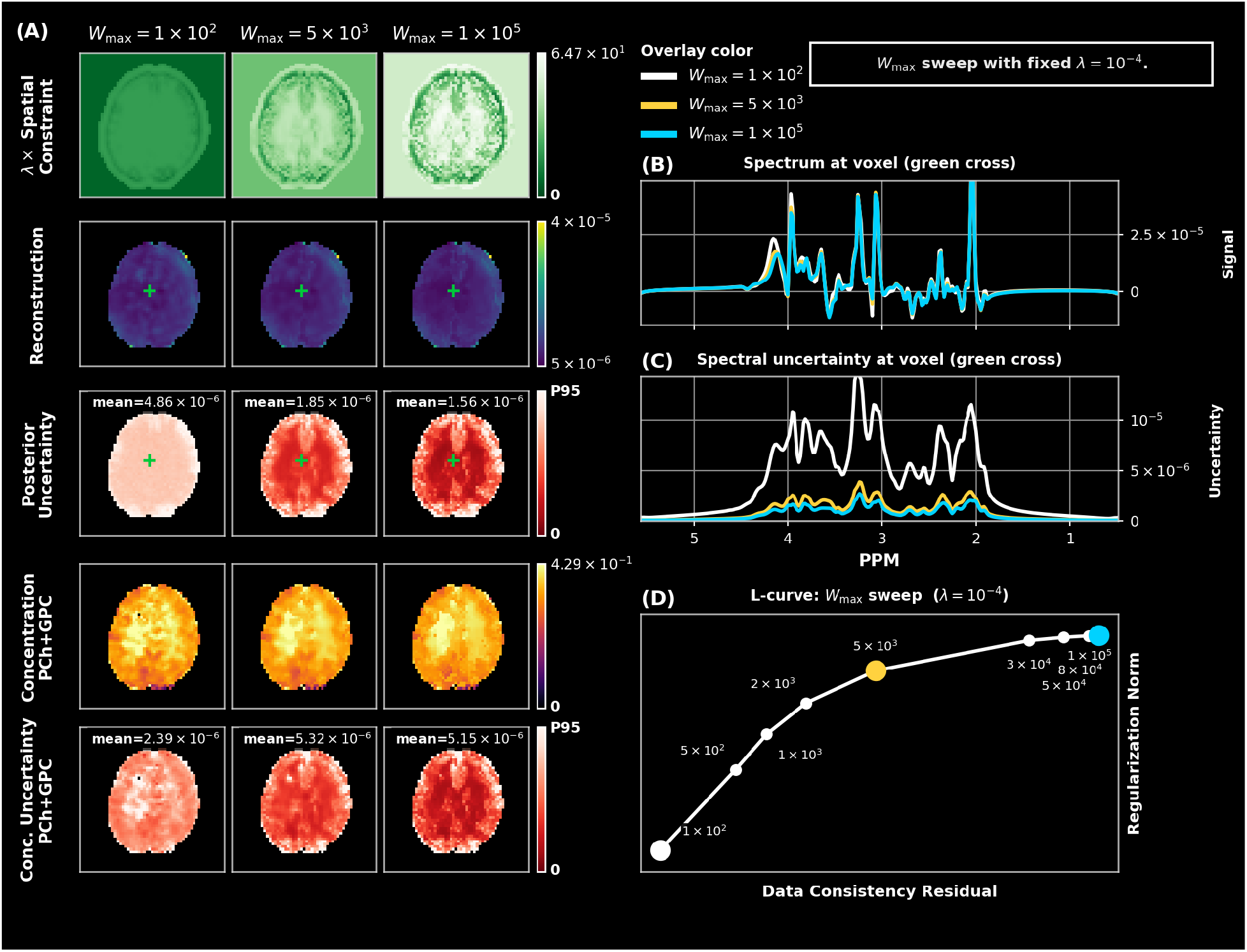
Effect of the edge-preserving constraint parameter *W*_max_ on SPICE reconstruction and posterior uncertainty. Representative results are shown for *W*_max_ = 1 × 10^2^, 5×10^3^, and 1×10^5^, with *λ* = 1×10^−4^ fixed. Left: spatial constraint, reconstruction, pre-fitting posterior uncertainty, PCh+GPC concentration, and concentration uncertainty. Right: reconstructed spectra, spectral uncertainty, and the L-curve across the *W*_max_ sweep. Increasing *W*_max_ produces more anatomically structured uncertainty while preserving tissue boundaries.

Pre-fitting posterior uncertainty changed from a relatively homogeneous distribution at low *W*_max_ to a more anatomically structured pattern at higher *W*_max_, with clearer tissue-level contrast (Fig. 5a). This effect was also observed in the metabolite concentration-uncertainty maps (Fig. 5a), which showed more spatially coherent within-tissue uncertainty. The L-curve changed most strongly at low *W*_max_ and progressively flattened at higher values, indicating a reduced effect of further increases in *W*_max_ (Fig. 5d).

### Effect of k-Space Undersampling

Fig. 6 shows the effect of retrospective k-space undersampling on SPICE reconstruction and the framework-quantified posterior uncertainty. With the central 30% of k-space fully retained, the SPICE reconstruction remained relatively stable for outer-k-space acceleration of *R* = 2, whereas *R* = 3 produced more noticeable spatial artefacts and signal attenuation in both the reconstructed maps and selected-voxel spectra.

**Figure 6.**
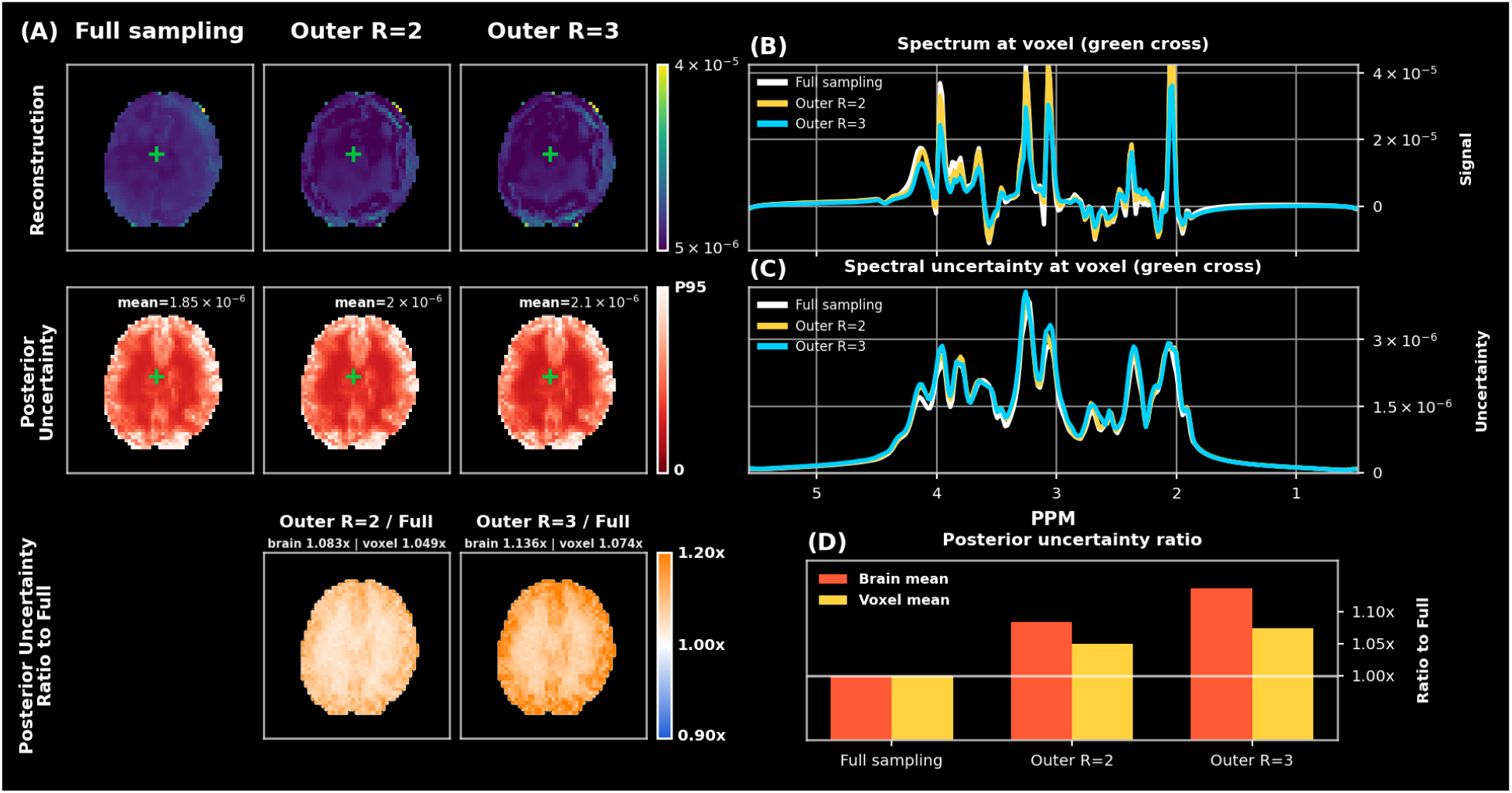
Effect of k-space undersampling on SPICE reconstruction and pre-fitting posterior uncertainty. Results are shown for the fully sampled acquisition and retrospective undersampling with outer-k-space acceleration factors of *R* = 2 and *R* = 3, while the central 30% of k-space is fully retained. The panels show the sampling patterns, SPICE reconstructions, posterior uncertainty, and reconstructed spectra and uncertainty spectra at the voxel indicated by the green cross. The lower panels show voxel-wise posterior-uncertainty ratios relative to the fully sampled reference and their summaries across the brain and at the selected voxel. Increasing undersampling produces greater reconstruction degradation and signal attenuation, while the overall spatial and spectral uncertainty patterns remain similar and their magnitude increases.

The spatial and spectral patterns of posterior uncertainty were largely preserved across the sampling schemes, while the uncertainty magnitude increased with greater undersampling. The voxel-wise uncertainty ratios were predominantly above unity for both *R* = 2 and *R* = 3, with a larger increase for *R* = 3, consistent with the whole-brain and representative-voxel summaries. Overall, stronger peripheral undersampling increased posterior uncertainty, although the increase remained modest relative to the reconstruction degradation observed at *R* = 3.

## Discussion

### Interpretation of Undersampling Uncertainty

In the undersampling experiment, the measured data **S** enter the gradient in Eq. 4 through a term independent of **U** and therefore disappear when forming the Hessian. Consequently, *H*(**Û**) depends on the sampling pattern through **F**^*H*^**F**, as well as on *B*_0_ modulation, the spectral basis, and spatial regularization, but not on the measured data realisation or reconstruction residual. This explains why structured artefacts such as ringing can appear in the reconstruction without producing equivalent patterns in the posterior uncertainty.

### Computational Considerations

In addition to voxel-wise conjugate-gradient estimation, we explored a truncated eigendecomposition of the reconstruction Hessian as an alternative acceleration strategy. Low-rank Hessian approximations have been widely used for large-scale Laplace uncertainty estimation [33–35]. Because contributions to H^−1^ are weighted by inverse eigenvalues, the smallest eigenpairs can be estimated using matrix-free LOBPCG [36]. The resulting global representation can be reused across voxels and is compatible with GPU computation, potentially reducing repeated voxel-wise solves.

### Generalisability to Other Low-Rank MRSI Reconstructions

Although the framework is developed for SPICE, the approach can be adapted to other differentiable low-rank MRSI reconstruction methods, including compartmentalised low-rank models [37] and low-rank tensor models [38]. For each method, the corresponding reconstruction Hessian can be derived and inverted approximately to obtain posterior covariance, which can then be propagated to reconstructed spectra or fitted quantities. The main method-specific component is therefore the reconstruction Hessian, while the covariance propagation and matrix-free computation remain applicable.

### Limitations and Future Work

The current framework primarily characterises posterior uncertainty arising from measurement noise under a fixed reconstruction model. It does not include uncertainty from motion, physiological fluctuations, or scanner drift. In addition, because the Hessian does not depend on the measured data realisation or reconstruction residual, systematic reconstruction bias caused by strong undersampling is not directly represented by the current uncertainty formulation.

Residual water and lipid contamination are not modelled explicitly, but residuals captured or entangled within the spectral basis can influence both pre-fitting uncertainty and, through spectral fitting, metabolite concentration uncertainty. This is consistent with the elevated pre-fitting and concentration uncertainty observed in peripheral voxels in Figs. 4 and 5, where both anatomical-boundary effects and residual water or lipid contamination may contribute. Extending the model to represent these sources of variability and bias explicitly would provide a more complete description of uncertainty.

The truncated LOBPCG approximation also requires further evaluation to determine its accuracy and an appropriate rank for practical use.

The current in vivo evaluation was performed in a small number of healthy participants. Further validation in larger datasets, different acquisition protocols, and patient populations will be required to assess the consistency of the framework-quantified uncertainty across different imaging conditions.

## Conclusions

We introduced a posterior uncertainty quantification framework for SPICE, which can be extended to more general regularized low-rank MRSI reconstruction. The proposed framework estimates posterior covariance from the inverse Hessian of the SPICE objective and propagates it to reconstructed spectra and metabolite concentrations. Matrix-free Hessian–vector products and conjugate-gradient solution of the voxel-wise linear systems avoid explicit construction or inversion of the full Hessian. Using repeated in vivo acquisitions, the quantified posterior uncertainty showed similar spatial and spectral patterns to empirical repeated-scan uncertainty, while the parameter experiments demonstrated how spatial regularization and k-space undersampling affect uncertainty. The framework can also be adapted to other differentiable low-rank MRSI reconstruction methods.

## Data Availability Statement

The source code used in this study is available at https://github.com/JasonLvernex/SPICE-P osterior-Uncertainty (commit SHA-1: `a216294584e360a184f72f92eb070777fd1d0f94`). The human MRSI data are not publicly available because public data sharing was not covered by the participant consent and ethical approval for this study. Data were acquired under a technical development protocol approved by local ethics and institutional committees.

## Acknowledgments

S.M.F. and W.T.C. jointly provided the primary supervision for this work and contributed equally as senior authors. S.J. contributed to the initial conceptual development of the study and provided senior guidance. This research was funded in whole or in part by the Wellcome Trust (225924/Z/22/Z) and supported by the NIHR Oxford Health Biomedical Research Centre (NIHR203316). The views expressed are those of the author(s) and not necessarily those of the NIHR or the Department of Health and Social Care. The Centre for Integrative Neuroimaging was supported by core funding from the Wellcome Trust (203139/Z/16/Z and 203139/A/16/Z). We thank the UK BBSRC (grant number BB/W019582/1) for support. T.L. is financially supported by the China Scholarship Council and the University of Oxford (202408270020). For the purpose of Open Access, the author has applied a CC BY public copyright licence to any Author Accepted Manuscript (AAM) version arising from this submission.

